# Glioblastoma stem cells show transcriptionally correlated spatial organization

**DOI:** 10.1101/2024.08.27.609918

**Authors:** Shamini Ayyadhury, Patty Sachamitr, Michelle M. Kushida, Nicole I Park, Fiona J. Coutinho, Owen Whitley, Panagiotis Prinos, Cheryl H. Arrowsmith, Peter B. Dirks, Trevor J. Pugh, Gary D. Bader

## Abstract

Glioblastoma (GBM) is an aggressive brain cancer with a poor survival rate. Despite hundreds of clinical trials, there is no effective targeted therapy. Glioblastoma stem cells (GSCs) are an important GBM model system. In culture, these cells form spatial structures that share morphological aspects with their source tumors. We collected 17,000 phase contrast images of 15 patient-derived GSC lines growing to confluence. We find that GSCs grow in characteristic multicellular patterns depending on their transcriptional state. Interpretable computer vision algorithms identified specific image features that predict transcriptional state across multiple cell confluency levels. This relationship will be useful in developing GSC screens where image features can be used to identify how GSC biology changes in response to perturbations simply by imaging cultured cells on plates.

## Introduction

Glioblastoma (GBM) is a brain cancer with poor overall survival despite extensive research and therapeutic testing^1,2^. GBM tumors and their component tumor-initiating glioblastoma stem cells (GSCs) are highly heterogenous, but follow characteristic patterns at the transcriptional level^3–5^. Since treatment outcomes are a function of these multicellular patterns, high-throughput therapeutic screening technology that models these could improve identification of targets and therapeutics^6–8^.

Organoids and two-dimensional (2D) cell culture models capture aspects of tissue-level cellular organization and are valuable in therapy development and preclinical testing^9^. An advantage of 2D cell culture systems is that they can be easily imaged, capturing interesting aspects of cellular organization, including cell shape, spatial relationships between cells and general properties of the space of all cells on the plate. They also ensure that every cell can be similarly influenced by media conditions, lessening heterogeneity related to artifacts of cellular clumps in culture. Traditional image-based therapeutic screening methods applied to cellular culture models generally extract few and relatively general phenotypes, such as growth rate or expression of select protein markers, from measured images^10^. Circumventing these limitations, recent image analysis advances support extracting rich, multidimensional information from single cell images^11,12^. However, most methods do not consider the organization of cells into multicellular structures and how these patterns relate to the human tissues modeled by 2D cultures.

To better understand the multicellular patterns present in GSC cultures, we analyzed 17,000 phase contrast microscopy images obtained from 15 patient-derived GSC cultures grown over 12-16 days. A set of 29 image analysis features, calculated based on image pixel distributions, was computed for each image. Unsupervised analysis reveals that these features naturally organize the GSCs along a spectrum that strongly correlates with a neurodevelopmental to injury response gradient and multiple brain cell type and inflammatory gene expression signatures^3,4,13–18^. Further, specific image features correlate with transcriptional states across multiple cell confluency levels. This relationship will be useful in developing improved GSC screens where we can identify how key aspects of GSC biology change in response to perturbations by imaging cultured cells on plates.

## Results

### GSCs show diverse morphometric and multicellular patterns in culture

To measure morphometry of 2D cell cultures, we used phase-contrast light microscopy to image GSC cultures from 15 patients, from day 1 until confluency (up to 16 days) at 4 or 12 hour intervals. This resulted in a database of 17,601 images (Supplementary fig 1A), previously used only for overall cell confluency measurements to develop controls in a chemical screen^19^. However, we observed strong pixel variation across time and confluency levels within these images, suggesting that the images contained additional information related to architecture (Supplementary fig 1B). Manual characterization of image patterns identified rich shape, texture, directionality and cell composition information within the images. Starting with single cells, we observed that some cells have homogenous texture (uniform or little pixel intensity variation) across a cell body or nucleus, making it difficult to ascertain the nuclear-cytoplasmic boundary (Figure 1A, top panel-white ✽). We observed broad cells, having multiple membranous protrusions with darkly pixelated structures resembling focal anchor points, attached to the well bottom (Figure 1A, top panel -yellow ✽✽), whereas other cell shapes are elongated in multiple ways (Figure 1A, top panel -red ✽✽✽) or exhibit angular, straight edged membranes with thin cytoplasm and contrasting darker nuclei (Figure 1A, bottom panel -red arrow). We also noticed smooth and rough textured nuclei (Figure 1A, top panel and bottom panel -white arrowhead, respectively). Some GSC cultures exhibit more than one of these morphometric features (e.g. G729) whereas others contain more limited cell shapes and textures (e.g. G797, G885, G895; Supplementary fig 1B). Next, we characterized visually prominent pattern variation present in images at higher cell confluency levels, which were accompanied by multicellular structure formation. We observed in some samples (G564; Figure 1B - left, G549, G799, G800; Supplementary fig 1B), cells have a tendency to align with their longitudinal axis against each other, exhibiting anisotropy, whereas in other samples we found isotropic patterning of cell clusters, where cell organization and grouping show no directionality (G523; Figure 1B -right, G566, G583, G729, G797, G837, G851, G861, G876, G885, G895; Supplementary fig 1B). We also observed differences in the spatial geometry of how cells relate to each other in space. For instance, cells from some patient samples (e.g. G837) organize themselves in a 2D layer, exhibiting efficient space utilization with little cell-cell overlap and no visible intercellular gaps (Figure 1C -top). This appears to be an inherent property of these cells and not due solely to confluency effects, as the property is visible at multiple confluency levels. However, cells from other samples, such as G876, grow in a largely overlapping manner, making extensive membrane projections that overlap nearby cells (Figure 1C -bottom). Overall, we observe a rich diversity of single cell morphology and emergent multicellular patterns in our cell culture images.

**Figure 1.**
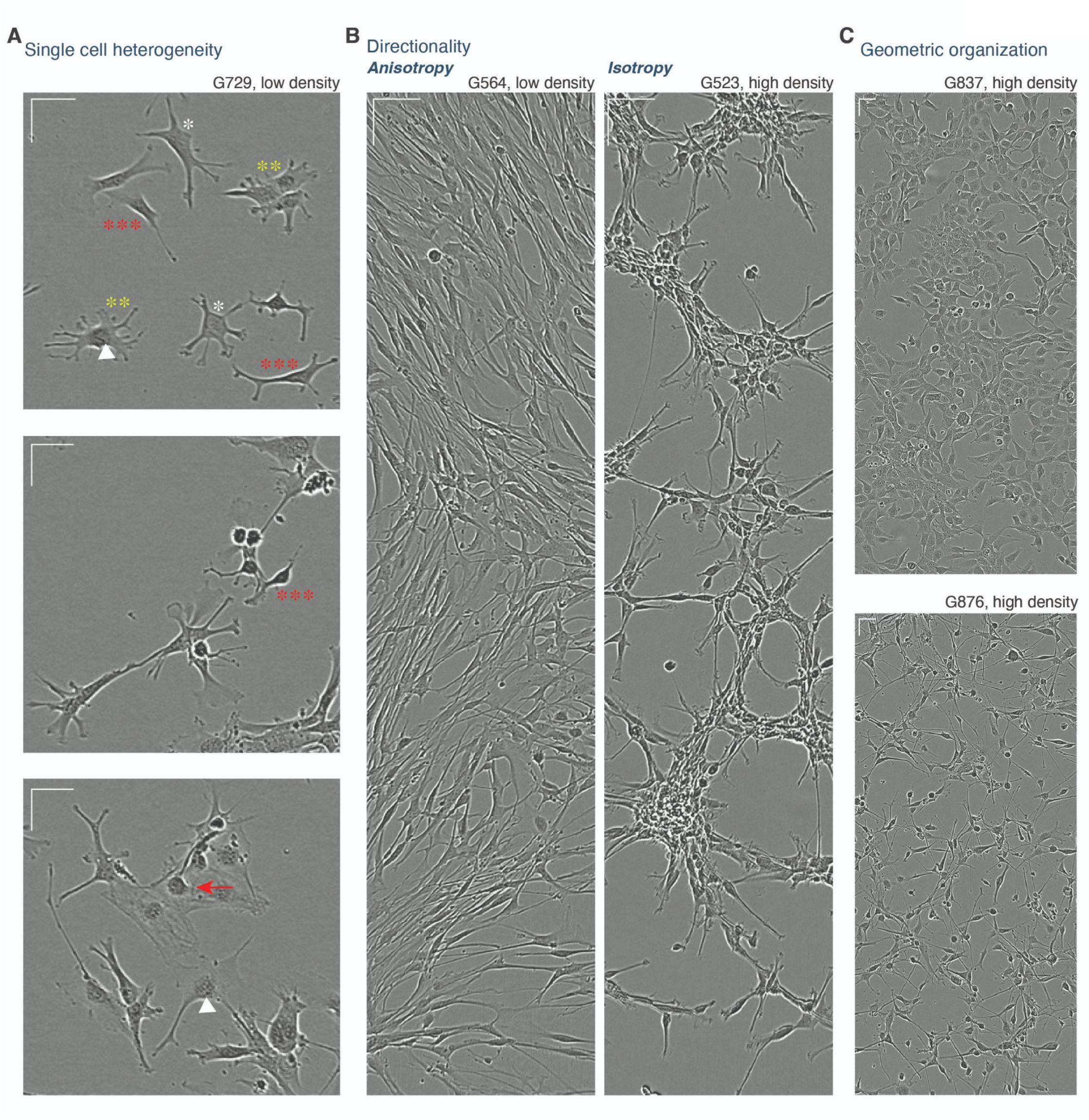
Diverse multicellular spatial patterning of patient-derived glioma stem-like cells in culture. A -C) Representative images of glioma stem-like cells exhibiting rich multicellular spatial patterning. *Scale representation = 50um for both x and y-axis bars*.

### Unsupervised factor analysis uncovers spatial patterns in GSC images that correlate with gene-expression signatures

We next asked if the multicellular patterns we observe in our images correlate with known biological programs, such as cell type or state gene expression signatures (Figure 2). To quantify the pattern variation at multiple confluencies, we used the CellProfiler software to compute a variety of statistical pixel distributional patterns (features) for each image. CellProfiler implements two classes of whole image feature extraction algorithms: gray level co-occurrence matrix (GLCM) and granularity spectrum (Supplementary Notes 1 and 2, materials and methods). Briefly, the GLCM is a symmetric frequency matrix derived from the raw pixel matrix of an image and represents the frequency distribution of pixel-pairs. From this frequency distribution, descriptive (i.e mean, variance, correlation) and spatial (i.e homogeneity, entropy, contrast) patterns of pixel-pair relationships across an image are calculated^20^. We also included the granularity spectrum, which captures the size range of structures in an image. This results in a total of 29 features representing pixel distribution patterns from an image.

**Figure 2.**
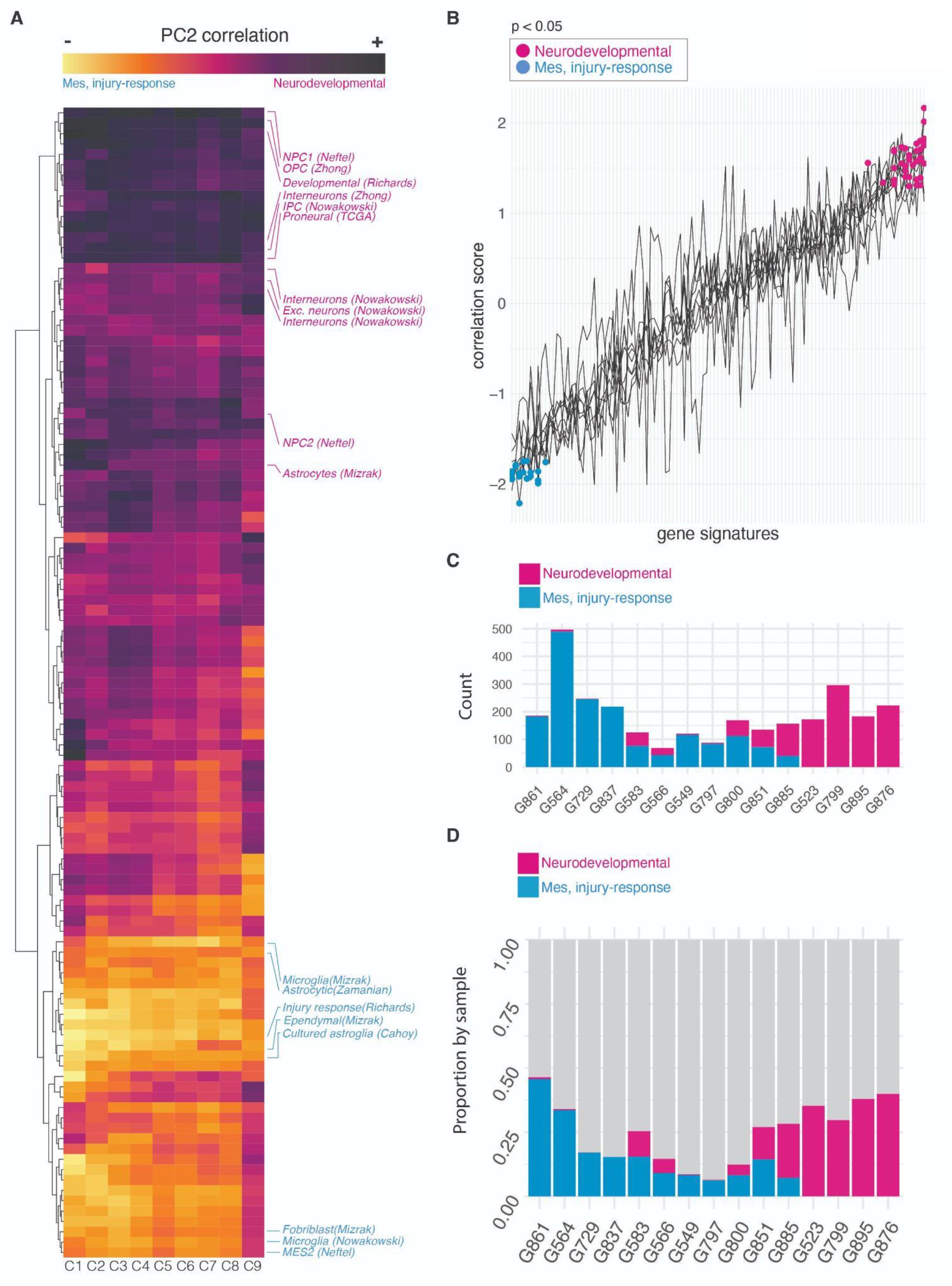
Image analysis features correlate with gene signatures. A) Heatmap of correlation scores between mean PC2 scores of samples from each confluency level and each of the 111 gene signature enrichment levels in transcriptomic data from matched samples. Select gene signatures are labeled and colored by their major biological category (magenta -neurodevelopmental, blue -mesenchymal-injury-response). B) The column vectors from heatmap in A), plotted as a line plot with statistically significant gene signatures colored by the major biological category they represent. C) The number of images from each sample contributing towards the top and bottom 25% of the PC2 component. D) The proportion of images by sample contributing towards the top and bottom 25% of PC2.

We analyzed these images over nine confluency levels to capture the changes in organizational geometries over varying cell densities (materials and methods, Supplementary fig 2A). We computed for each image a feature vector using the 29 image features described above. We normalized these feature vectors for images separately for each confluency level, to control for cell density, and then performed principal component analysis (PCA) over each confluency group. We found that PC1 and PC2 carried the bulk of the variance (39-59% total, across confluency levels) (Supplementary fig 2B). We next asked if the PCs correlate with known biological signals. To answer this, we analyzed bulk RNA-seq datasets matched with our 15 GSC samples. We scored each RNA-seq sample using 111 gene signatures from human fetal and adult brain cell types and GBM tissues from a range of studies (Supplementary table 1, materials and methods^3,4,13–15,17,18,21^). To correlate our image feature vectors by image, and our 111 gene signature vectors by sample, we averaged our image feature vectors by sample for each principal component and confluency level (materials and methods, Supplementary fig 2C). We then computed the Pearson correlation coefficient for each averaged component with each of the 111 gene signatures (Supplementary fig 2D). Both PC1 and PC2 showed biological correlation across confluency levels, though PC2 gave a more generalizable association with gene expression (Figure 2A-B, Supplementary fig 3). PC2 is correlated with signatures representing OPCs, NSCs, neurons and astrocytes, and anti-correlated with signatures representing microglia, mesenchymal and injury response phenotypes (Figure 2, Supplementary table 2). Thus, GSCs in culture exhibit biologically meaningful spatial patterning with neurodevelopmental and mesenchymal/injury-response biological programs showing distinct geometric patterns well separated in PC space.

### The GSC multicellular spatial pattern to gene expression correlation is maintained across all growth phases, but relevant image features change over time

We next interpreted the image features correlated with the neurodevelopmental and mesenchymal/injury-response transcriptional gradient. Overall, these image features show broad patterns that change with cell density (Supplementary fig 4). For instance, granularity spectrum values increase as cell density increases (Supplementary fig 4A). This trend is expected as at lower cellular densities there are more single or smaller clusters of cells. The GLCM derived image features show a more complex relationship with confluency levels, with some features positively or negatively correlated with cell density and others showing a non-linear relationship (Supplementary fig 4B-C).

Image features also captured an additional axis of variation, distributing images along the neurodevelopmental-mesenchymal/injury-response gradient (PC2) (Figure 3). Despite maintaining a consistent overall correlation with the neurodevelopmental and mesenchymal/injury-response gradient across confluency levels, the image features driving PC2 variance change over cell growth phases, from low to mid to high confluency (Figure 3A). For instance, small granularity features are consistently associated with lower cell densities. However, within these lower cell densities, the PC2 loadings are higher for the low granularity features with respect to the neurodevelopmental GSC culture images, indicating smaller structures within these images, in contrast to images from the mesenchymal/injury response GSC culture images of similar densities (Figure 3A, Supplementary fig 5A, Supplementary table 5). In contrast, PC2 loadings for the mid granularity features are stronger in the low density mesenchymal/injury response GSC culture images (Figure 3A, Supplementary fig 5B, Supplementary table 5). GLCM-derived features also show a clear biological association with the neurodevelopmental-mesenchymal/injury-response gradient. For instance, the informational measure features (IMC1 and IMC2), both measuring the mutual information along the horizontal and vertical image axes (Supplementary Note 1), are the strongest PC2 feature loadings for the neurodevelopmental images (Figure 3A). IMC1 is strongly correlated with the neurodevelopmental signatures at the low-mid confluency levels whereas IMC2 exhibits a stronger correlation with the neurodevelopmental samples at high confluency levels (Figure 3B, Supplementary table 5). This reflects the symmetry of multicellular structures at two different density ranges and shows how neurodevelopmental cell states grow in more symmetric multicellular patterns than mesenchymal/injury-response cell states. Another example is the inverse difference moment, which measures homogeneity by penalizing high contrast areas. It is associated with neurodevelopmental images at mid-confluency levels (Figure 3A and C, Supplementary table 5). The correlation feature, measuring similarities in pixel neighborhood patterns in the image, is also more strongly associated with neurodevelopmental images at mid confluencies (Figure 3D, Supplementary table 5). Differences in pixel-pair values along different axes in different directions can be attributed to variation in the sharp boundaries or the nature of pixel-intensity shifts that form as cells meet each other along their membranes, overlap cellular processes or exhibit anisotropy, forming variable local contrast patterns. The data indicates a more consistent cell population patterning within the neurodevelopmental images compared to the mesenchymal and injury-response samples and this may relate to consistent overall cellular composition, resulting in cells maintaining similar textures even as they grow into larger cellular networks. Overall, image analysis features highlight organizational differences along the neurodevelopmental to mesenchymal/injury-response biological gradient that are preserved despite the changing spatial organization patterns of cells in culture during growth.

**Figure 3.**
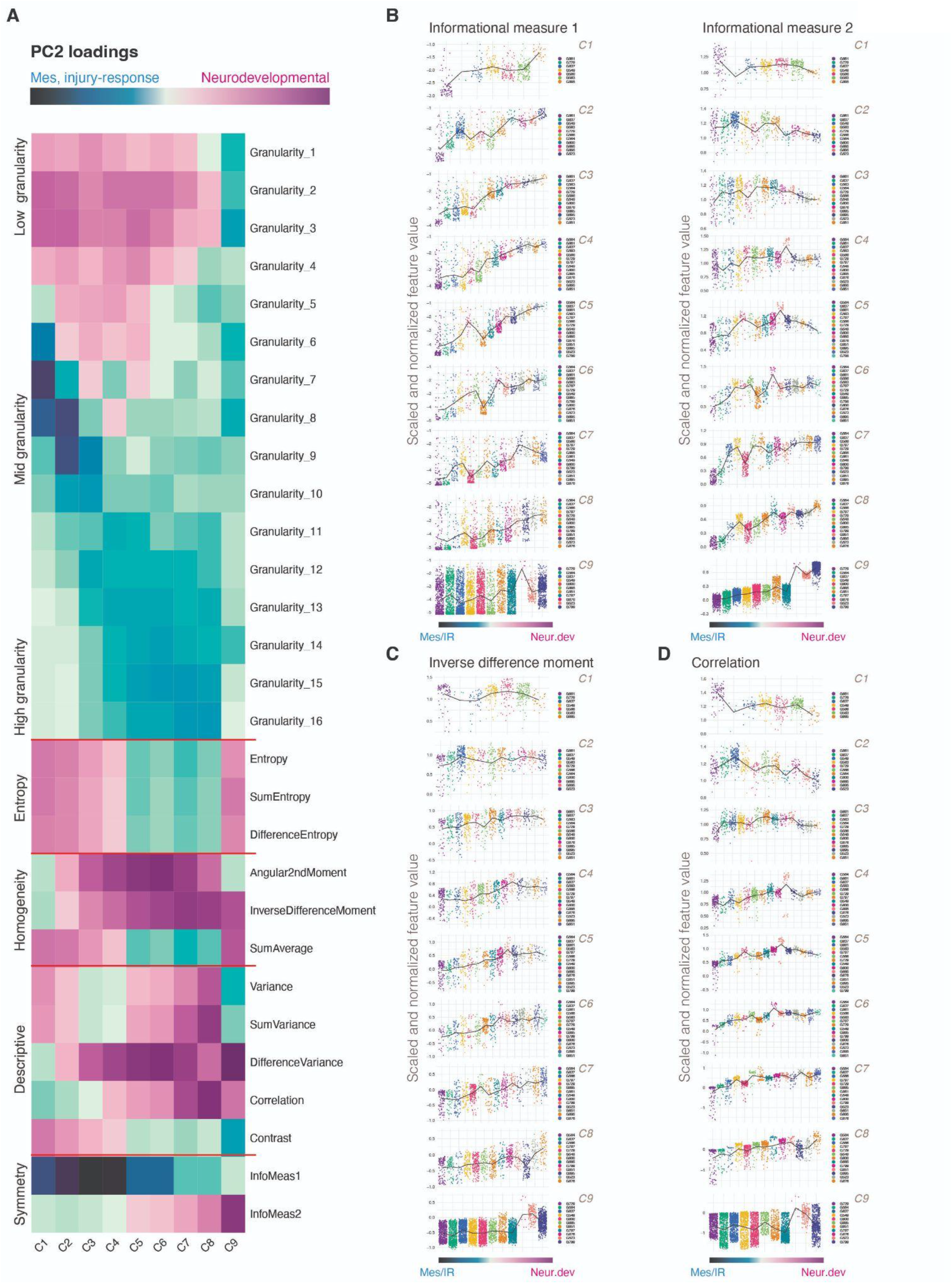
Identifying image analysis features that best correlate with GSC neurodevelopmental/injury response transcriptomic gradient. A) A heatmap of the PC2 loadings for all 29 image features used in this study. B-D) Representative distribution plots of images grouped by samples and ordered by their mean PC2 (correlated with the GSC transcriptional gradient), with a line-plot showing the trend. Pearson correlation coefficients and corresponding p-values shown were obtained by correlating image feature values and image PC2 component scores for all images from a confluency group. X-axis represents samples ordered by their mean PC2 value and y-axis represents the scaled and normalized feature value. ** represents p value < 0.01 and *** represents p value < 0.001.

## Discussion

Microscopic images of cells growing over time on 2D plates are a rich source of biological information, but are traditionally used to extract single values, such as cellular growth rate, for analysis. The multi-cellular spatial patterns visible in these images are defined by cell-type orientation, space utilization, steric hindrances between cells and cell shape. Here we show that these patterns vary along temporal and biological axes. In particular, cultured glioblastoma stem cells form patterns that strongly correlate with a transcriptomic gradient expressing neurodevelopmental pathways on one side and mesenchymal and injury-response/inflammation pathways on the other. Mesenchymal/injury response GSCs show increasingly more diverse multicellular patterns while neurodevelopmental samples exhibit more stable patterning as they grow. We also find that spatial patterning varies with time and yet maintains a strong correlation with the GSC transcriptomics gradient strongly suggesting that as spatial structures evolve they are constrained by underlying biological programs.

A key open question is how single cell morphometric features relate to the higher-order multicellular patterns we studied. Is there more spatial pattern information available at the multicellular level than can be discerned from studying individual cells? Unfortunately, we couldn’t address this as we had relatively few single cell examples to study in our data. Further, extracting high-quality single cell shapes from multicellular neighborhoods is challenging for various reasons, such as cellular overlap and difficulty resolving extended cellular membrane protrusions from each other, even when using expert manual annotation. Improved cell segmentation and imaging methods, including those that consider more detailed time series to resolve overlapping cells, will be required to extract enough single cell images from multicellular structures in 2D culture images to learn these multiscale relationships^22^.

Extending our 2D study to other models such as live and fixed tissue slices imaged with diverse microscopy technologies, as well as three-dimensional systems, such as organoids, is expected to provide richer information than can be extracted from the brightfield phase-contrast images we analyze here. However, our analysis approach considers whole images including all structures in view, thus should be generally compatible with other imaging systems and models, only requiring image feature adaptation to cover additional image-based information as needed.

Ultimately, we hope that further study of relationships between images and functional genomics data will identify gene expression programs that are correlated with specific multicellular structures, extending our interpretability capability to identify causal relationships. This will be useful to improve our understanding of disease models and develop new therapies that consider the architecture of growing multicellular systems.

## Materials and Methods

### Glioblastoma stem cell culture

Fresh tumor samples were obtained from patients during operative procedures following informed consent. All experimental procedures were performed in accordance with the Research Ethics Board at The Hospital for Sick Children (REB1000025582, REB0020010404), the University Health Network, the University of Calgary Ethics Review Board and the Health Research Ethics Board of Alberta, Cancer Committee and Arnie Charbonneau Cancer Institute Research Ethics Board (REB HREBA-CC-160762)^4,19^. GSC samples were derived as previously described (See Richards et al. supplementary note 1^4^). Only images from adherently grown cultures were used in this study.

### Phase contrast imaging

The phase contrast image data were collected as established^19^. Briefly, two thousand GSCs, maintained between passages 8-12, were plated adherently in 384-well CELLBIND plates (Corning) and imaged using the Incucyte Zoom™ live cell imaging system (Essen Biosciences). Cells were imaged with a Nikon 10x objective using phase-contrast mode every 4 – 8 hours until the experimental end point of a confluent plate (12-16 days). Culture media was changed every 5 days. Images used in this study were from wells grown untreated (stem cell media only). Original image size was at 1392 × 1040, with a pixel size of 1.22um/pixel, before it was exported out at scaled size of 1266 × 944.

### Frame extraction before image analysis

Images were saved from the microscope as time-lapse videos (.mp4 video files) and openCV was used to extract image frames from those videos. Each phase contrast image was saved as a Tag Image File Format (TIFF) file.

### Whole Image analysis

- Mask generation with Ilastik

For each TIFF image, we used the *ilastik* software (pixel classification module) to generate background segmentation masks that exclude any ‘acellular’ regions of the plate/image during downstream computation^23^ (Supplementary fig 1A). Every second image until t=24 hours and every 6th image thereafter until t=168 hours were used for training. Each sample dataset was processed independently with its own training run, to account for sample-level batch effects. Two classifiers, background and foreground were created within each Ilastik session to separate plate pixels from cell pixels. A single colored stroke, with stroke width ranging from 1-3, was used to mark the background/plate area and a single stroke with a different color was used to mark the cell occupied regions. Any debris present was included as part of the foreground. This process was repeated for a sampling of images (every 3rd-5th image). If necessary, training was continued using additional unused images. Each image used in training only received 1-2 training strokes per classifier group per image. After training, the *ilastik* pixel classifier for each sample dataset was run on a computer cluster using all images. The background and cell masks generated were used downstream for image data extraction using *Cellprofiler*’s built-in image analysis algorithms.

- Image pre-processing and image analysis

First, raw images were screened using *Cellprofiler’*s *‘MeasureImageQuality’* module to remove poor quality images^20^. After trying out several of the module’s parameters, we found that the *‘PowerLogLogSlope’* (PLLS) was sufficient to remove aberrant images^24^. The images removed by PLLS were manually verified to ensure that they were blurry or corrupted by noticeable artifacts (e.g. pipette plastic objects, well bottom scratches) and no exceptions were found. The foreground masks, representing cellular regions, were used to calculate the area of cells occupying each image (Supplementary fig 2A). Following this, the *‘MeasureTexture’* and *‘MeasureGranularity’* modules were used to extract gray-level co-occurrence matrix pixel pair distributions and the granular or size distribution patterns from these images using 29 image analysis features, with default settings (Supplementary Notes 1 and 2). The *‘MeasureGranularity’* algorithm uses 16 different pixel sizes of openings/closings or structuring elements as its default. These were used to analyze the granularity of each image and its relative size. The *‘MeasureTexture’* module measures the Haralick features which consists of 13 features, measuring various aspects of pixel-pair spatial distribution patterns, using the gray scale co-occurrence matrix. For the Haralick features, a scale factor from 0-3 was used. The scale factor refers to the directionality of the GLCM matrices (‘north’, ‘south’, ‘east’, ‘west’ and explained in Supplementary note 1). These four directions perfectly correlate with each other, thus we included only a single scale (direction=‘east’) from our data analysis (scale = 0 in CellProfiler).

- Normalization

After computing image features, normalization was carried out by sample by converting each feature value into a within-sample fraction. The data matrix is composed of rows representing individual images and columns representing one of the 29 image features. Each feature vector or column was multiplied by a diagonal matrix, where the diagonal represented the scaling factor to transform each feature vector as a fraction of the maximum feature value.

I_c_ = diag(a_1_, a_2_, a_3_…., a_f_) [*f*_1_,*f*_2_,*f*_3_, …, *f*_n_]^T^

I = Image matrix, c = image column, a_f_ = maximum value of pixel feature in each column c, *f*_*n*_= feature vector

Following this, z-normalization was applied by row to obtain a normalized feature vector for each image.

- Batch correction and data binning into nine confluency levels

Principal component analysis (PCA) over all images is strongly driven by the time-point the images were taken at and the sample (batch effect). To mitigate batch effects, we grouped the images by time-point, to ensure that images from the same plate and time-point were grouped together. Next we formed batches of images based on the mean total cell area per image while grouped by time-point for each sample set to form different confluency levels. Altogether, we separated the images into 9 confluency levels (Supplementary fig 2A).

- Principal component analysis of whole image analysis.

PCAtools version 2.8.0 with default parameters was used for PCA dimensional reduction of all images.

### Bulk RNA-seq analysis

Bulk RNA sequencing data from Richards et al was used^4^. Gene set variation analysis (GSVA)^25^ scores previously computed were used for this study using previously published gene sets (Supplementary table 4). Briefly, GSCs, maintained between 8-12 passages, were grown till 70-80% confluency before being harvested for bulk RNA sequencing. 15 bulk RNAseq datasets, matched by sample with the imaging datasets, were annotated using gene lists from various published studies to mark for neural-progenitor, neuronal, oligodendrocyte progenitor, oligodendrocyte, mesenchymal, astrocytic, immune and tumor phenotypes including the injury-response/developmental signature^3,4,13–15,17,18,21^. *The original gene signature scoring pipeline will be made available upon publication of this manuscript*. Downregulated gene signatures were removed from the analysis as these formed mostly redundant pairs with upregulated signatures, resulting in 111 gene signatures used in this study.

### Pearson correlation analysis between whole image and bulk RNA-seq datasets

For each confluency group, we used Pearson correlation analysis to compute the correlation coefficients between the normalized image features and the 111 gene signatures obtained from bulk RNAseq. First, the mean PC scores were calculated for each GSC for all 15 samples, resulting in a vector of 15 mean PC scores which were then correlated against each of the 111 gene signature vectors, where each vector comprises the GSVA scores for a single unique signature across 15 GSCs (Supplementary fig 2D). We changed the directions of all anti-correlated pairs to ensure a common direction so that we could compare biologically meaningful associations across confluency levels and gene signatures.

To obtain the top and bottom 25% images along PC1 and PC2, first the PC1 and PC2 scores for all images were z-normalized by confluency levels. Next, z-normalized matrices were concatenated to form a single dataframe and ranked by PC2 scores. The top 25% and bottom 25% ranked images were subsetted to calculate the proportion of sample contribution. P values obtained from the corr.test output were considered significant when values fell below 0.01. The correlation coefficients and p-values can be found in Supplementary table 5.

## Supporting information

supplementary figures and notes (excluding supp tables)

## Dataset repositories and code availability

*All code, CellProfiler pipelines, and ilastik pipelines used in the manuscript will be made available on Github and Zenodo upon acceptance of publication*.

## Authorship

SA conceived the study. SA, TJP and GDB designed the study with methodological development by SA and GDB. PS, MMK, NIP and FJC designed, developed and executed the experimental work for cell culture and imaging under the supervision of PP, CHA and PBD. All computational analysis was done by SA, except for the bulk RNA sequencing pipeline from OW, under the supervision of GDB. The manuscript was written by SA, TJP and GDB. The manuscript underwent final revisions with input from all authors.

## Notes

### Competing Interest Statement

The authors have declared no competing interest.

## References

1. Stupp, R. et al. Radiotherapy plus concomitant and adjuvant temozolomide for glioblastoma. N. Engl. J. Med. 352, 987–996 (2005).

2. Louis, D. N. et al. The 2021 WHO Classification of Tumors of the Central Nervous System: a summary. Neuro Oncol. 23, 1231–1251 (2021).

3. Neftel, C. et al. An integrative model of cellular states, plasticity, and genetics for glioblastoma. Cell 178, 835-849.e21 (2019).

4. Richards, L. M. et al. Gradient of Developmental and Injury Response transcriptional states defines functional vulnerabilities underpinning glioblastoma heterogeneity. Nat. Cancer 2, 157–173 (2021).

5. Couturier, C. P. et al. Single-cell RNA-seq reveals that glioblastoma recapitulates a normal neurodevelopmental hierarchy. Nat. Commun. 11, 3406 (2020).

6. Ayensa-Jiménez, J. et al. Mathematical formulation and parametric analysis of in vitro cell models in microfluidic devices: application to different stages of glioblastoma evolution. Sci. Rep. 10, 21193 (2020).

7. Albanese, A. et al. Multiscale 3D phenotyping of human cerebral organoids. Sci. Rep. 10, 21487 (2020).

8. Kelley, M. E. et al. High-content microscopy reveals a morphological signature of bortezomib resistance. eLife 12, (2023).

9. Ryoo, H., Kimmel, H., Rondo, E. & Underhill, G. H. Advances in high throughput cell culture technologies for therapeutic screening and biological discovery applications. Bioeng. Transl. Med. (2023) doi:10.1002/btm2.10627.

10. Ayestaran, I. et al. Identification of Intrinsic Drug Resistance and Its Biomarkers in High-Throughput Pharmacogenomic and CRISPR Screens. Patterns (N Y) 1, 100065 (2020).

11. Rohban, M. H. et al. Virtual screening for small-molecule pathway regulators by image-profile matching. Cell Syst. 13, 724-736.e9 (2022).

12. Kobayashi, H. et al. Label-free detection of cellular drug responses by high-throughput bright-field imaging and machine learning. Sci. Rep. 7, 12454 (2017).

13. Verhaak, R. G. W. et al. Integrated genomic analysis identifies clinically relevant subtypes of glioblastoma characterized by abnormalities in PDGFRA, IDH1, EGFR, and NF1. Cancer Cell 17, 98–110 (2010).

14. Zhong, S. et al. A single-cell RNA-seq survey of the developmental landscape of the human prefrontal cortex. Nature 555, 524–528 (2018).

15. Cahoy, J. D. et al. A transcriptome database for astrocytes, neurons, and oligodendrocytes: a new resource for understanding brain development and function. J. Neurosci. 28, 264–278 (2008).

16. Liddelow, S. A. et al. Neurotoxic reactive astrocytes are induced by activated microglia. Nature 541, 481–487 (2017).

17. Mizrak, D. et al. Single-Cell Analysis of Regional Differences in Adult V-SVZ Neural Stem Cell Lineages. Cell Rep. 26, 394-406.e5 (2019).

18. Nowakowski, T. J. et al. Spatiotemporal gene expression trajectories reveal developmental hierarchies of the human cortex. Science 358, 1318–1323 (2017).

19. Sachamitr, P. et al. PRMT5 inhibition disrupts splicing and stemness in glioblastoma. Nat. Commun. 12, 979 (2021).

20. Bray, M.-A. & Carpenter, A. E. Quality Control for High-Throughput Imaging Experiments Using Machine Learning in Cellprofiler. Methods Mol. Biol. 1683, 89–112 (2018).

21. Zamanian, J. L. et al. Genomic analysis of reactive astrogliosis. J. Neurosci. 32, 6391–6410 (2012).

22. Lange, M. et al. Zebrahub - Multimodal Zebrafish Developmental Atlas Reveals the State Transition Dynamics of Late Vertebrate Pluripotent Axial Progenitors. BioRxiv (2023) doi:10.1101/2023.03.06.531398.

23. Berg, S. et al. ilastik: interactive machine learning for (bio)image analysis. Nat. Methods 16, 1226–1232 (2019).

24. Bray, M.-A., Fraser, A. N., Hasaka, T. P. & Carpenter, A. E. Workflow and metrics for image quality control in large-scale high-content screens. J. Biomol. Screen. 17, 266–274 (2012).

25. Hänzelmann, S., Castelo, R. & Guinney, J. GSVA: gene set variation analysis for microarray and RNA-seq data. BMC Bioinformatics 14, 7 (2013).

