## supplementary figures and notes (excluding supp tables) for "Glioblastoma stem cells show transcriptionally correlated spatial organization"

### SUPPLEMENTARY FIGURES 1-5

#### Glioblastoma stem cells show transcriptionally correlated spatial organization

Shamini Ayyadhury[1,2,14], Patty Sachamitr[3,4], Michelle M. Kushida[3], Nicole I Park[3] , Fiona J. Coutinho[3], Owen Whitley[1], Panagiotis Prinos[4], Cheryl H. Arrowsmith[2,4,5], Peter B. Dirks[3,6,7,8,9], Trevor J. Pugh[2,5,10,14], Gary D. Bader[1,2,6,11,12,13,14]

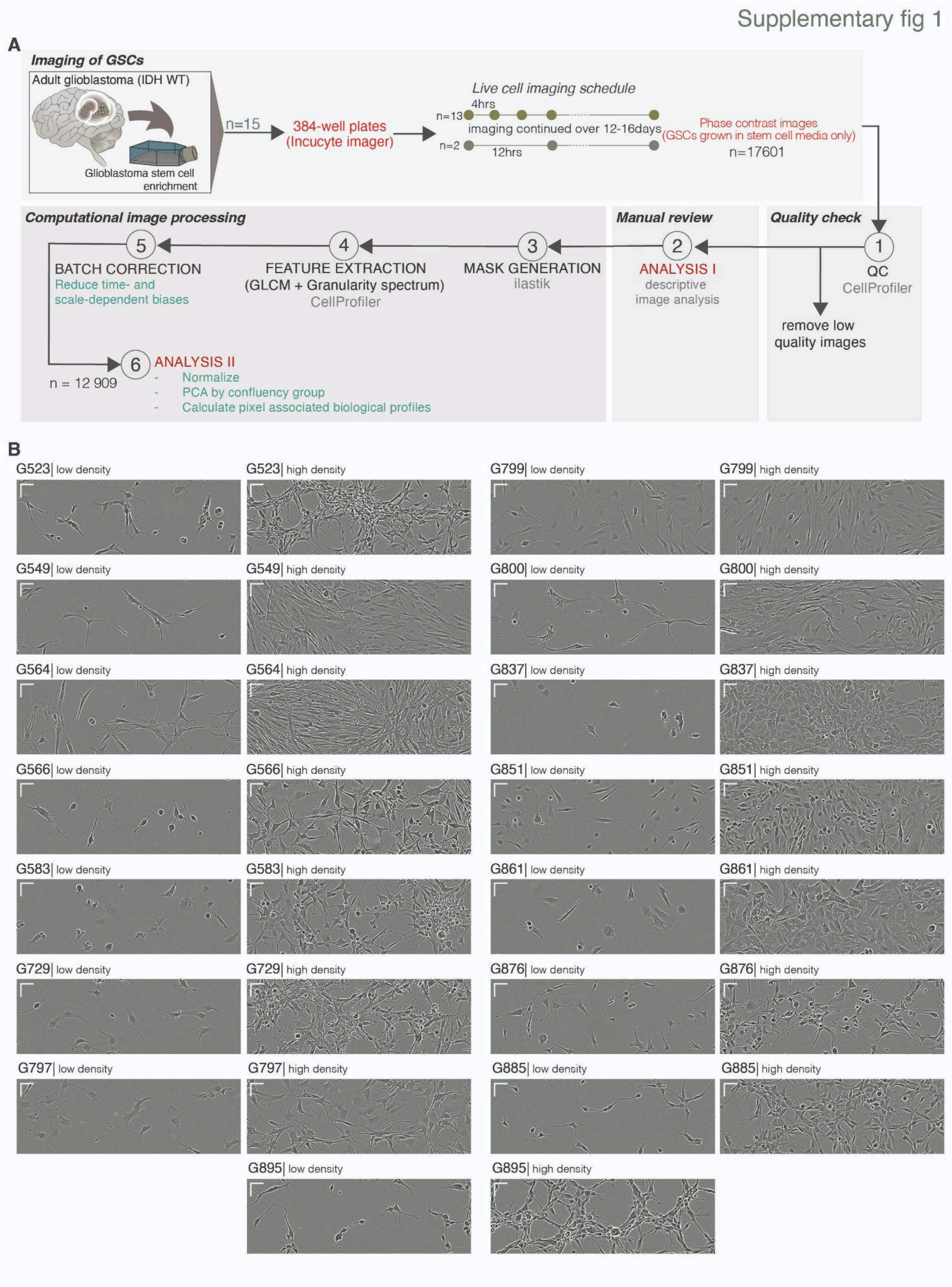


**Supplementary fig 1. Analysis workflow and representative images from our 15 GSC samples.** A) Workflow for manual and computational analysis and B) representative images from 15 patient-derived GSCs from low and high-density phase-contrast images. *Scale representation = 50um for both x and y-axis bars.*

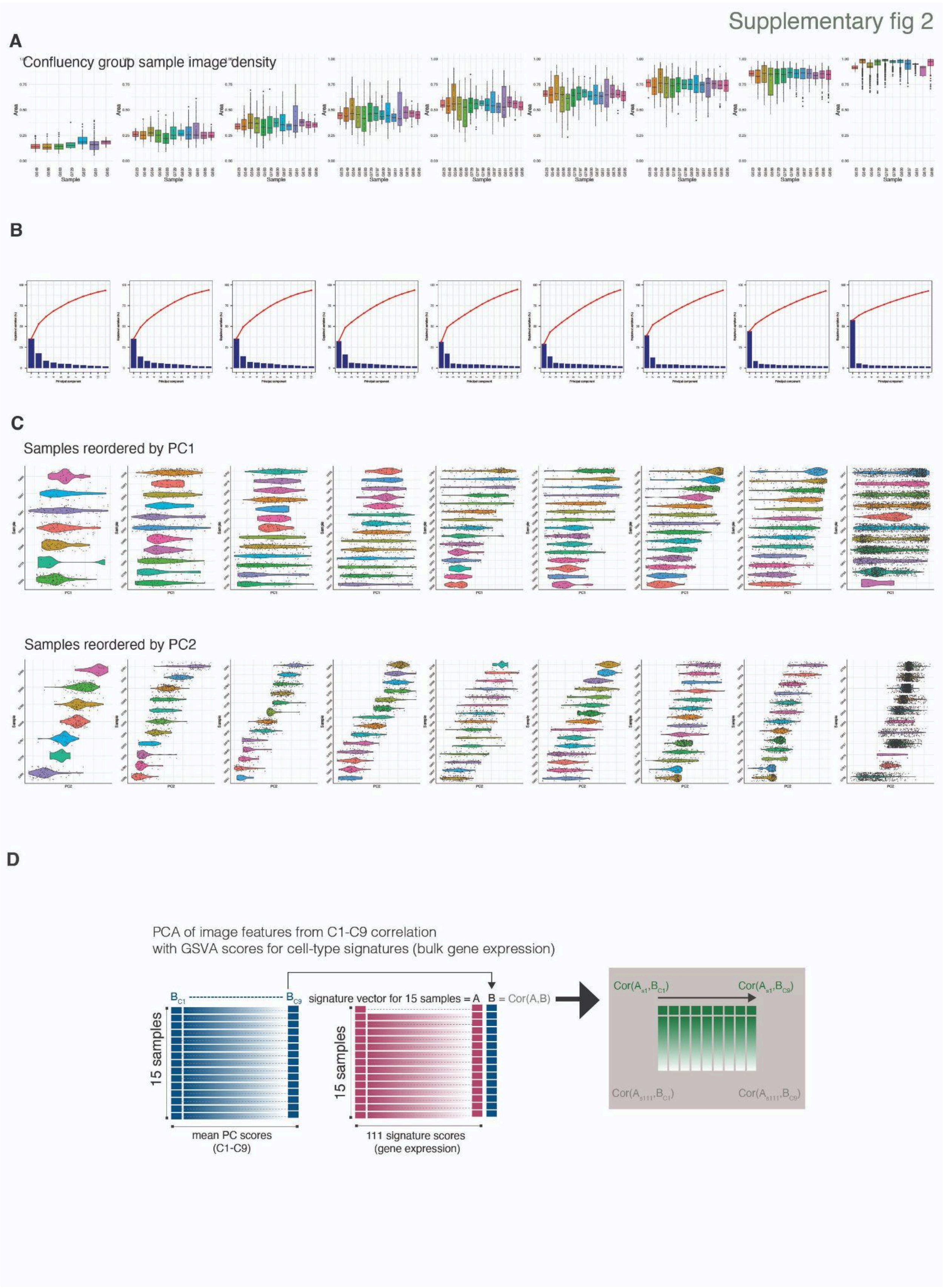


**Supplementary fig 2. Definition of nine confluency levels.** A) Nine confluency levels and the cell density variation for all images across samples for each one. B) Principal component cumulative variance for nine confluency levels. C) Mean sample PC2 scores shown for all confluency levels, with samples in each level ordered by the sample mean PC2. D) Schematic of correlation analysis performed between PCA of image features and gene signatures from bulk RNA expression after PC directional correction (see materials and methods) The correlation between the mean sample PC scores and each of the 111 gene signatures (s1-s111) was computed for all nine confluency groups (c1-c9).

Supplementary fig 3

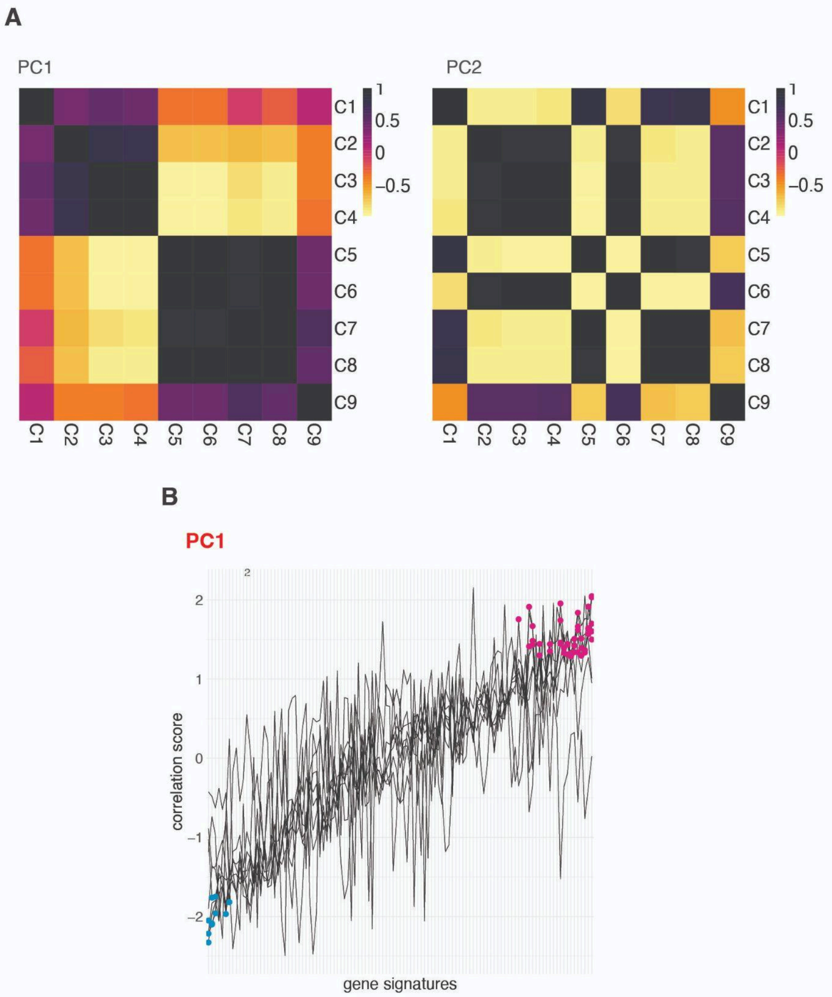


**Supplementary fig 3. Evaluating PC1 and PC2 relationship with gene signature scores.** A) Correlation matrices between confluency levels for PC1 and PC2 (after directional correction). B) PC1 correlations with 111 gene signatures for all nine confluency levels.

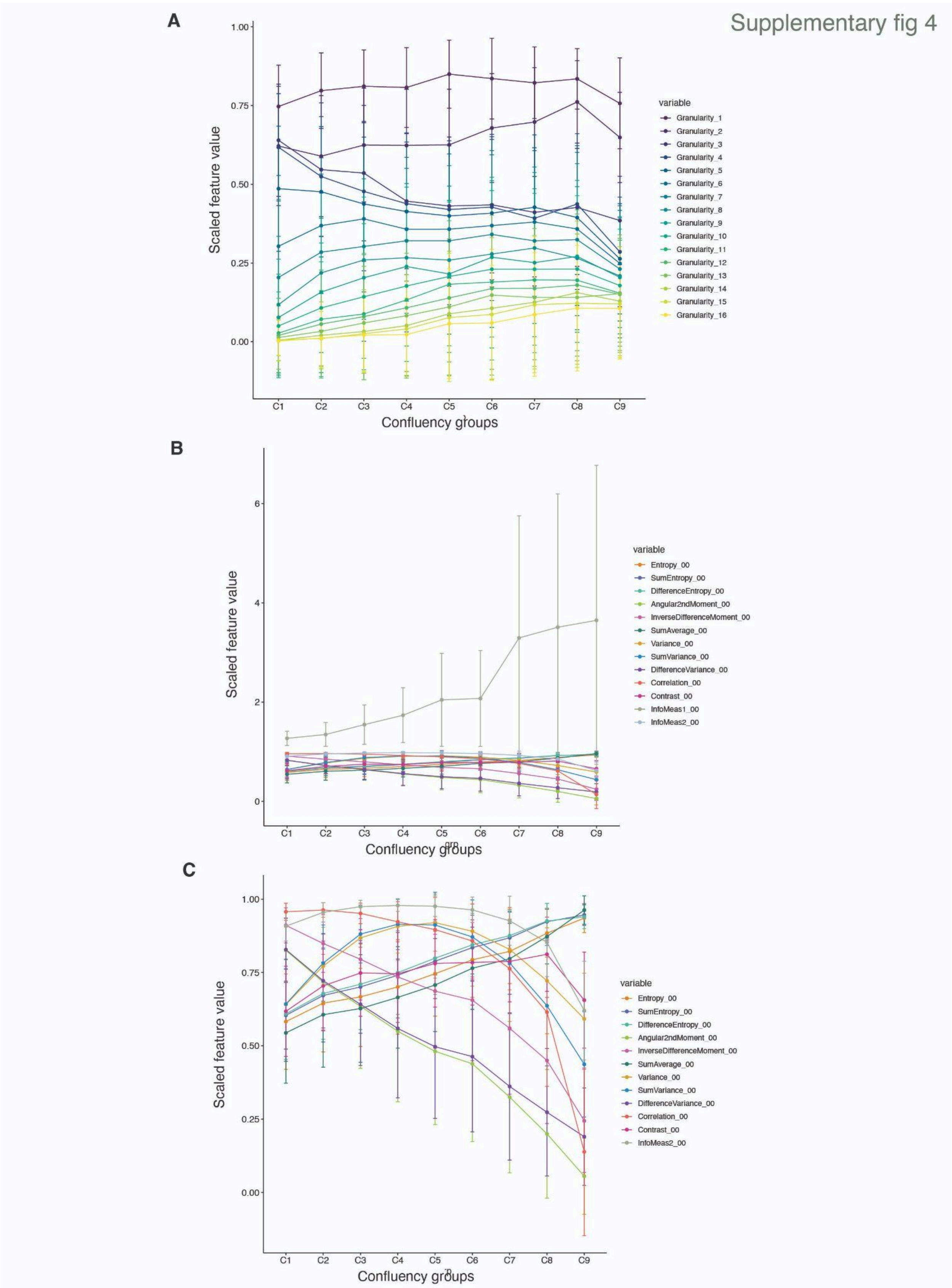


**Supplementary fig 4. Image features show broad patterns of variation along cell density levels.** A) The variation in granularity spectrum across different cell densities. Y-axis is the scaled image feature value obtained by dividing the feature vector with the maximum feature value**.** B) GLCM features vary along confluency levels, with features belonging to the same or similar families following the same pattern. Y-axis is the scaled image feature value obtained by dividing each feature vector by the maximum feature value. C) Same as B) but after removing Informational Measure 1 as an outlier.

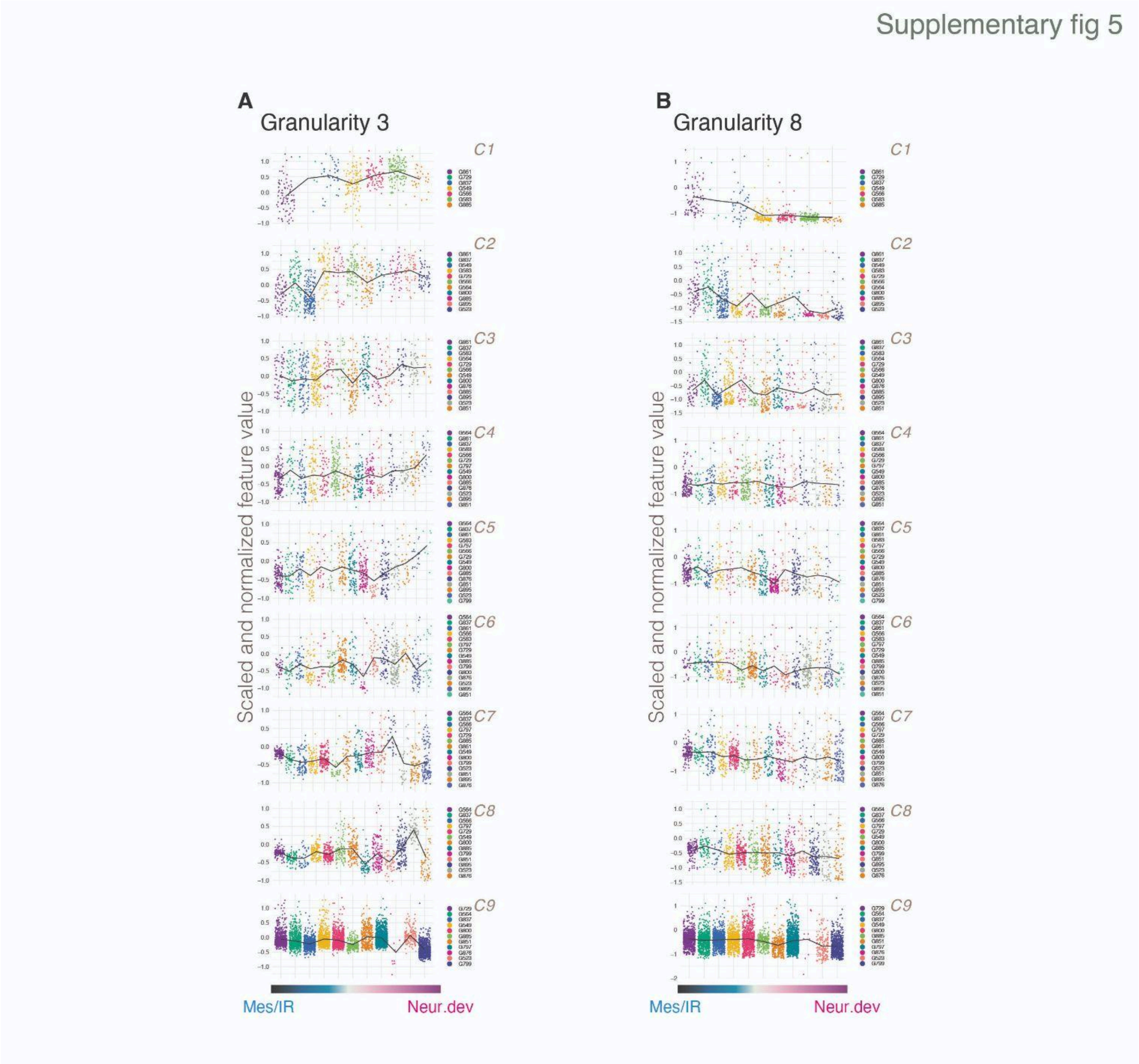


**Supplementary fig 5. Representative image features correlated with GSC gradient.** A) Granularity 3 shows a stronger correlation with the GSC gradient at lower confluency levels, with a higher correlation with the neurodevelopmental groups, indicating smaller structural units or patches in these images whereas B) mesenchymal/injury response images showed larger patches or clusters of cells at lower densities as indicated by the higher signals from the mid granularity spectrum (shown here using a representative example of Granularity 8).
** represents p value < 0.01 and *** represents p value < 0.001.

### SUPPLEMENTARY NOTES 1-2

#### Glioblastoma stem cells show transcriptionally correlated spatial organization

Shamini Ayyadhury[1,2,14], Patty Sachamitr[3,4], Michelle M. Kushida[3], Nicole I Park[3] , Fiona J. Coutinho[3], Owen Whitley[1], Panagiotis Prinos[4], Cheryl H. Arrowsmith[2,4,5], Peter B. Dirks[3,6,7,8,9], Trevor J. Pugh[2,5,10,14], Gary D. Bader[1,2,6,11,12,13,14]

**Supplementary Note 1**

#### A. Understanding GLCM computation

##### a. Toy Example Image

Each image is defined by a matrix of pixels. The range of each pixel is defined by the bit size. (This is in contrast to the gray level co-occurrence matrix which is symmetric and the rows and columns are defined by the pixel bit value (i.e 8 bit = 256 rows/columns))

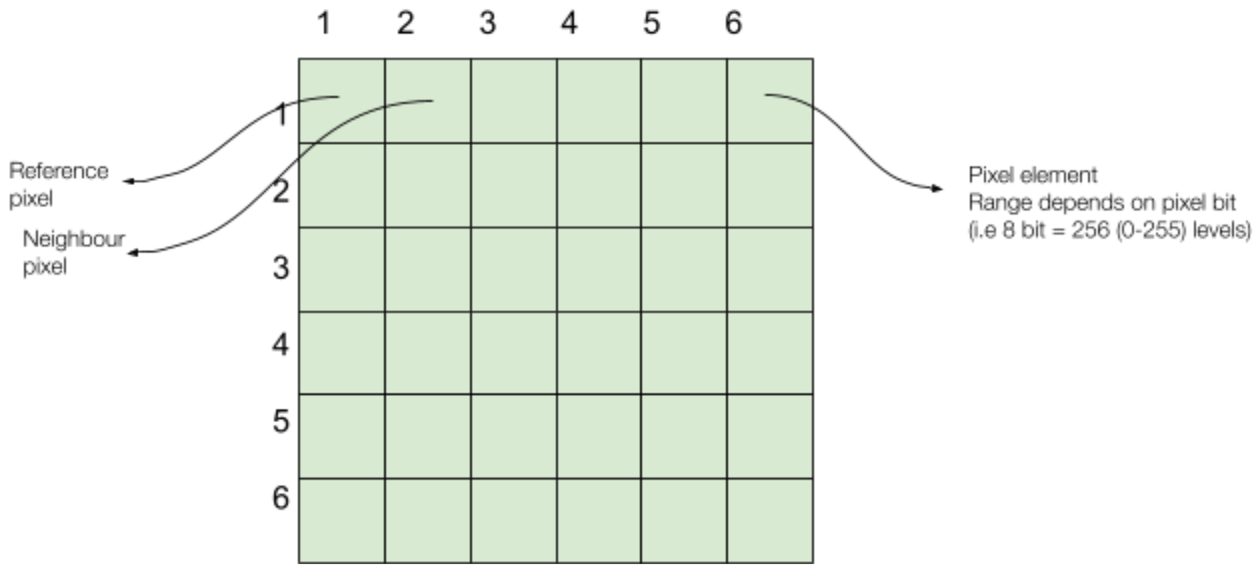


**Supplementary note fig 1.** *6 pixel x 6 pixel.*

##### b. Construction of the GLCM

The figure below will explain step by step the construction of the GLCM and the subsequent mathematical derivatives.

**Figure 1A.** The GLCM model can be constructed from using the entire image or a window can be designated to analyze a portion of an image. Here we generate a gray level co-occurrence matrix for a window (15 x 15).

**Figure 1B.** This is the 15 x 15 image pixel matrix.

**Figure 1C**. First a 256 x 256 matrix is constructed. The dimensions are determined by the pixel bit and in this example we are analyzing a 8 bit image which generates 256 gray levels.
Starting from the left and moving towards the right, the pixel on the left serves as a reference pixel and the pixel on the right the neighbor pixel. Once reaching the rightmost column, the process is repeated from right to left, where the right pixel serves as the reference pixel and the left pixel serves as the neighbor pixel.
In this case, the original 15 x 15 image window has only 1 (133,135) pixel pair in the forward (left to right) direction but 2(133,135) pixel pairs in the backward (right to left) direction. We tabulate the counts for these pixel pairs.

**Figure 1D**. We obtain the final horizontal matrix by adding the forward (A) and backward (B) matrices. Note that $B = A^T$, therefore we can simplify this step by computing only the counts matrix for one direction and adding the transpose to it.

GLCM is constructed in 4 main directions - horizontal, vertical and 2 diagonal. We show the horizontal construction as an example.

**Figure 1E.** The matrix is divided by the total number of counts calculated to derive a probability distribution matrix which we call the normalized GLCM.

**Figure 1F.** This probability matrix is used to evaluate the spatial pixel distribution patterns found in an image.

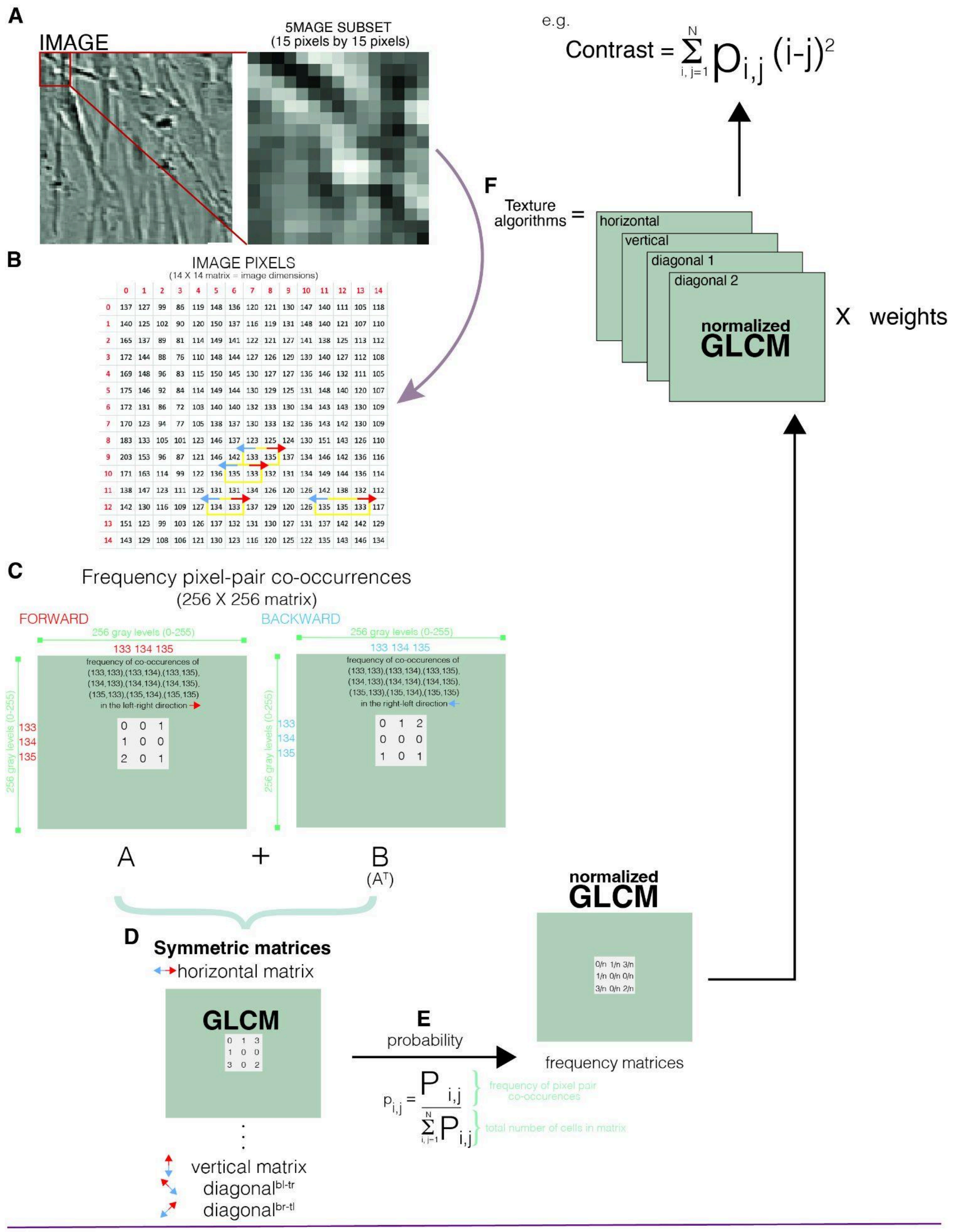


**Supplementary Note figure 2. GLCM model.** This figure shows how the gray level co-occurence matrix is constructed using a test image segment weighing 15 x 15 pixels.

##### c. GLCM can be constructed using different scale factors

Reference-Neighbour pixel pairs can take on different range or scales as shown below

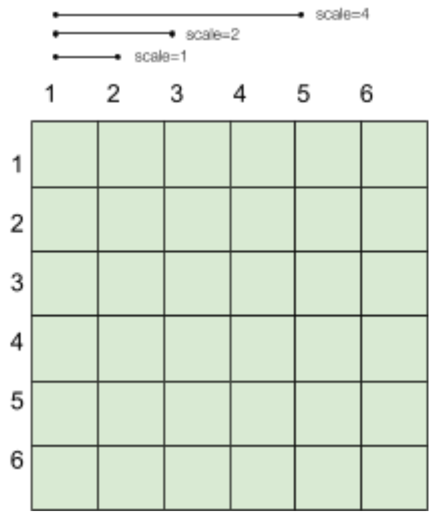


**Supplementary note fig 3.** Scales in GLCM construction. For scale =1, the reference pixel and neighbor pixel are adjacent, whereas for scale=2, the neighbor pixel is 2 pixels away.

For our example below, we will use scale=1.

##### d. GLCM-derived mathematical algorithms

**Legend**

$Ng \ = \ gray\ levels\ (pixel\ bit)$

The scale factor determines the distance between the reference and neighbor pixels that's used to compute the GLCM.

$$p_x(i) \ = \ ith\ entry\ in\ the\ marginal\ probability\ matrix \ = \ \sum_{i=1}^{Ng} p(i,j)$$

$$p_y(j) \ = \ jth\ entry\ in\ the\ marginal\ probability\ matrix \ = \ \sum_{j=1}^{Ng} p(i,j)$$

$$p_{x+y}(i + j) \ = \ probability\ distribution\ of\ the\ sum\ of\ two\ gray\ levels \ = \ \sum_{i=0}^{Ng}\sum_{j=0}^{Ng} p(i,j)$$

$i + j \ \in [0, 1,..., 1Ng]$ $\quad x\ \&\ y\ represent\ gray\ levels\ not\ coordinates$

$$p_{x-y}(i + j) \ = \ probability\ distribution\ of\ the\ difference\ of\ two\ gray\ levels \ = \ \sum_{i=0}^{Ng}\sum_{j=0}^{Ng} p(i,j)$$

$|i - j| \ \in [0, 1,..., Ng]$ $\quad x\ \&\ y\ represent\ gray\ levels\ not\ coordinates$

$\mu_x,\ \mu_y,\ \sigma_x, \sigma_y\ are\ the\ main\ standard\ deviations\ of\ p_x\ and\ p_y$

***Gray level co-occurrence matrix is constructed from the pixels of images from the normalized GLCM.***

*GLCM (normalized)*

$$p(i,j) = \frac{P(i,j)}{\sum_{i=1}^{Ng}\sum_{j=1}^{Ng} P(i,j)}$$

**GLCM derived hand-engineered pixel features**

1. *Contrast*
   Computes a larger weight when the difference between the reference and neighbour pixel is higher and hence results in a larger output when the image is dominated by sharp variations in pixel values.
   $$\sum_{i=1}^{Ng}\sum_{j=1}^{Ng} (i - j)^2 p(i,j)$$

2. *Inverse difference moment*
   Measures the homogeneity of an image window. The higher the value, the more homogeneous the gray level pixel-pairs are in the area inspected. Here pixel pairs moving away from the diagonal (and hence having more contrast) reduce the homogeneity value and vice versa.
   $$\sum_{i=1}^{Ng}\sum_{j=1}^{Ng} \frac{p(i,j)}{1 + (i-j)2}$$

3. *Angular Second Moment (ASM)*
   The ASM is a measure of the homogeneity as well, except that it is simply the square of the normalized GLCM and effectively gives equal weight to all pixel pair differences as opposed to IDM where pixel pairs with larger contrast are penalized more.
   $$\sum_{i=1}^{Ng}\sum_{j=1}^{Ng} p(i,j)^2$$

4. *Sum of squares (Variance)*
   Computes the variance in the reference pixels and their weighted contribution towards the GLCM frequency distribution and sums it across all gray level pixel pairs. A higher value means the presence of higher contribution of pixel pairs with larger pixel value differences. Related to Contrast.
   $$\sum_{i=1}^{Ng}\sum_{j=1}^{Ng} (i - \mu)^2 p(i,j)$$

5. *Sum Variance*
   *Variance of sum of gray levels*
   $$\sum_{i=2}^{2(Ng)} (i - Sum\ Average)^2 p_{x+y}(i)$$

6. *Difference Variance*
Variance of difference of gray levels

$$\sum_{i=1}^{Ng} (i - \mu)^2 p_{x-y}(i)$$

7. *Entropy*
Measure of randomness of intensity values in an image

$$-\sum_{i=1}^{Ng}\sum_{j=1}^{Ng} p(i,j)\, log(p(i,j))$$

8. *Sum Entropy*
Measure of randomness of sum of gray level intensity values in an image

$$-\sum_{i=1}^{2(Ng)} p_{x+y}(i)\, log(p_{x+y}(i))$$

9. *Difference Entropy*
Measure of randomness of difference in gray level intensity values in an image

$$-\sum_{i=1}^{Ng} p_{x-y}(i)\, log(p_{x-y}(i))$$

10. *Sum Average*
*The average sum of gray levels*

$$\sum_{i=2}^{2Ng} ip_{x+y}(i)$$

11. *Correlation*
The correlation between gray-level pixel pairs from a GLCM.

$$\sum_{i=1}^{Ng}\sum_{j=1}^{Ng} \frac{(i \bullet j)p(i,j) - \mu_x\mu_y}{\sigma_x\sigma_y}$$

12. *Information measures of correlation (IMC)*
IMC1

$$\frac{Entropy - HXY1}{max(HX, HY)}$$

$$HX = -\sum_{i}^{Ng} p_x(i)log(p_x(i)) = entropy\ of\ p_x$$

$$HY = -\sum_{j}^{Ng} p_y(j)log(p_y(j)) = entropy\ of\ p_y$$

$$HXY1 = -\sum_{i}^{Ng}\sum_{j}^{Ng} p(i,j)log[p_x(i)p_y(j)]$$

IMC2

$$[1 - exp(- 2(HXY2 - Entropy)]^{1/2}$$

$$HXY2 = - \sum_{i}^{Ng} \sum_{j}^{Ng} p_x(i)p_y(j)log[p_x(i)p_y(j)]$$

#### Supplementary Note 2

##### Granularity spectrum

Granulometry computes the size distribution by using a series of morphological operators (functions). The granularity spectrum used in this study uses a range of structuring elements varying in width from 1 pixel to 16 pixels. Each structuring element applied over the image will remove pixels that are equal or smaller than itself. The differences between the proportion of pixels removed in successive structuring element applications produces the size distribution inherent in the image (Figure 1).

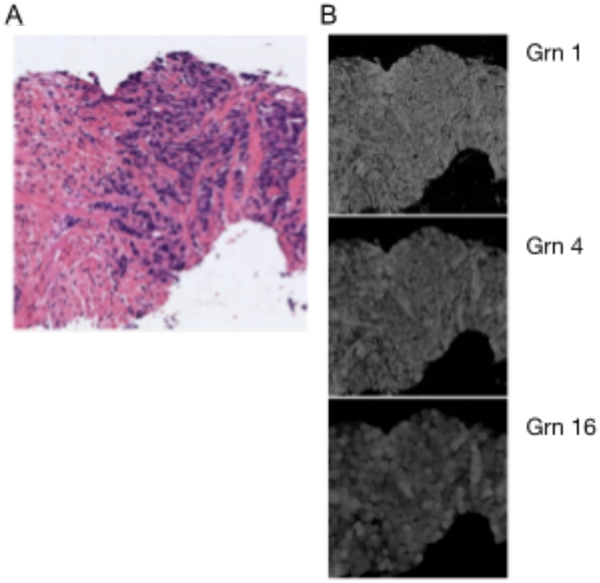


**Figure 1.** A) IHC image of prostate cancer tissues. B) Granularity profiles for IHC image on A). Increasing structuring element diameter results in identification of larger structures from the image. *Adapted from Esteban et al.*[1]

References

1. Esteban, Á. E. *et al.* A new optical density granulometry-based descriptor for the classification of prostate histological images using shallow and deep Gaussian processes. *Comput. Methods Programs Biomed.* **178**, 303–317 (2019).
